# Applicability of AlphaFold2 in the modeling of dimeric, trimeric, and tetrameric coiled-coil domains

**DOI:** 10.1101/2024.03.07.583852

**Authors:** Rafal Madaj, Mikel Martinez-Goikoetxea, Kamil Kaminski, Jan Ludwiczak, Stanislaw Dunin-Horkawicz

## Abstract

Coiled coils are a common protein structural motif involved in cellular functions ranging from mediating protein-protein interactions to facilitating processes such as signal transduction or regulation of gene expression. They are formed by two or more alpha helices that wind around a central axis to form a buried hydrophobic core. Various forms of coiled-coil bundles have been reported, each characterized by the number, orientation, and degree of winding of the constituent helices. This variability is underpinned by short sequence repeats that form coiled coils and whose properties determine both their overall topology and the local geometry of the hydrophobic core. The strikingly repetitive sequence has enabled the development of accurate sequence-based coiled-coil prediction methods; however, the modeling of coiled-coil domains remains a challenging task. In this work, we evaluated the accuracy of AlphaFold2 in modeling coiled-coil domains, both in modeling local geometry and in predicting global topological properties. Furthermore, we show that the prediction of the oligomeric state of coiled-coil bundles can be achieved by using the internal representations of AlphaFold2, with a performance better than any previous state-of-the-art method (code available at https://github.com/labstructbioinf/dc2_oligo).

## Introduction

Coiled coils are protein structural motifs consisting of two or more α-helices, oriented parallel or antiparallel, that wind around a central axis to form rod-like bundles (Lupas et al., 2017; Woolfson, 2023). This winding, also known as supercoiling, is associated with the periodic interlocking of side chains localized in the hydrophobic core of the bundle. The basis of this interlocking, an interaction known as knobs-into-holes, is considered to be the hallmark of coiled coils. It places the side chain of a core residue (knob) of one helix into a cavity (hole) formed by residues of the opposite helix. At the sequence level, coiled-coil domains are typically formed by 7-residue repeats (heptad) in which residues are labeled from a through g. Residues at two positions, a and d, are typically hydrophobic and face the bundle axis, forming the hydrophobic core. The difference between the α-helix periodicity, which is fixed at approximately 3.62 residues per turn (Szczepaniak et al., 2021), and the coiled-coil sequence periodicity of 3.5 (corresponding to 2 core positions per 7 heptad residues) is accommodated by the supercoiling of the coiled-coil structure and the formation of left-handed bundles (Figure 1A).

**Figure 1.**
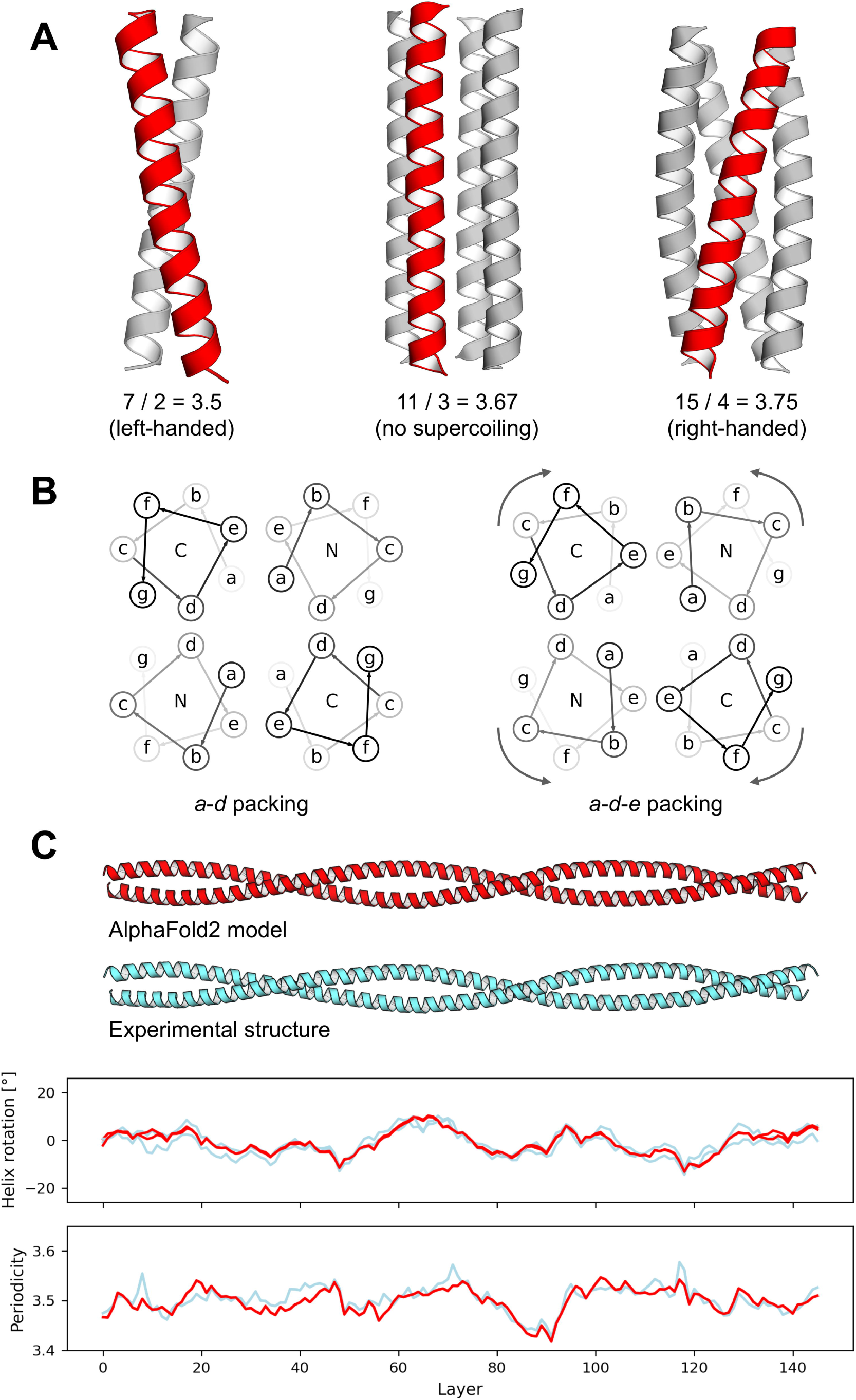
Coiled-coil domains and quantification of their structures. (A) Examples of coiled-coil structures with different periodicities. The most typical periodicity is 7/2, which induces the left-handed twist of a bundle, while others such as 11/3 or 15/4 are considered non-canonical. (B) Helical wheel diagrams of antiparallel coiled coils with heptad positions labeled a-g. Letters in the center of the helices indicate whether they are viewed from the N- or C-terminal side. The left diagram corresponds to the canonical a-d geometry, while the right diagram corresponds to the extended a-d-e geometry caused by a counterclockwise axial rotation of all helices by about 26 degrees, viewed from the N-terminus. (C) Experimental structure and corresponding AlphaFold2 model of the tropomyosin fragment. The results of the measurement of two structural parameters, helix axial rotation and periodicity, are shown below in scale. These parameters are defined separately for each individual layer of an input structure (layer is defined as a set of n residues, where n corresponds to the number of helices in the bundle, roughly localized on a plane perpendicular to the bundle axis).

Although heptads represent the vast majority of known coiled-coil domains, deviations from this canonical periodicity are possible and have been observed, very often as local non-canonical repeats interspersed between arrays of heptads, but also as global non-heptad coiled-coil domains (Martinez-Goikoetxea and Lupas, 2023). For example, one of the most common non-heptad repeats is the hendecad (Figure 1A), which is 11 residues long and contributes three residues to the hydrophobic core. Since the average spacing between core residues (11/3=3.67) is very close to the periodicity of an alpha helix (3.62), coiled-coil bundles based on hendecad repeats show almost no supercoiling. Unlike hendecads, pentadecads are 15 residues long, contribute four residues to the core (15/4=3.75), and cause right-handed supercoiling (3.75>3.63). These departures from the heptad periodicity are necessarily accompanied by the partial loss of the knobs-into-holes interactions, as some of them become knobs-to-knobs, with the side chains of the core residues (knobs) pointing toward the central axis instead of toward a hole. The canonical knobs-into-holes packing can also be disrupted by axial rotation of the constituent helices, which does not alter supercoiling, but can lead to the formation of knobs-to-knobs interactions. For example, counterclockwise rotation of helices (viewed from their N-terminal ends) in an antiparallel 4-helix bundle of 7/2 periodicity by about 26° relative to the canonical packing results in the formation of an a-d-e core, with residues in positions a pointing toward the center of the bundle and residues in positions d and e flanking them (Figure 1B).

Due to their regular nature, coiled-coil structures can be fully described by parametric equations using tools such as ISAMBARD, CCCP, Twister, and SamCC (Wood et al., 2017; Grigoryan and Degrado, 2011; Strelkov and Burkhard, 2003; Szczepaniak et al., 2021; Crick, 1953). Such descriptions allow structural parameters to be traced down to single-residue resolution, highlighting subtle differences within and between coiled-coil domains (Figure 1C). The most important parameters describing coiled-coil structures are their topology, i.e., the number and relative arrangement of helices (parallel vs. antiparallel), the degree of supercoiling (from left-handed to nearly straight to right-handed), and the axial rotation of helices, which defines the architecture of the hydrophobic core. Such a detailed view is important for relating the structural properties of coiled-coil domains to the function of the proteins that contain them. For example, the structural parameters of the HAMP domain, a small signaling 4-helix coiled coil, have been shown to be tightly coupled to the enzymatic activity of downstream domains (Ferris et al., 2011).

The high frequency with which coiled-coil domains are found (Szczepaniak et al., 2021) as well as their well-understood sequence-structure relationship have motivated the development of many computational tools for their detection and modeling. Tools for sequence annotation of coiled-coil domains, such as COILS (Lupas et al., 1991) and DeepCoil2 (Ludwiczak et al., 2019), exhibit remarkably robust performance, which can be attributed to the structural constraints that cause even unrelated coiled-coil sequences to share similar patterns. Despite our ability to predict coiled coils from sequence, their structural modeling remains a challenging problem that has been addressed by programs such as CCFold (Guzenko and Strelkov, 2018) and Rosetta (Das et al., 2009), which are based on fragment modeling, and ISAMBARD (Wood et al., 2017), CCBuilder (Wood and Woolfson, 2018), BeammotifCC (Offer et al., 2002), and CCCP (Grigoryan and Degrado, 2011), which use the aforementioned parametric equations. However, none of these programs is universally applicable. Some are limited to certain oligomeric states or require additional data besides a sequence, while others can only model perfectly regular bundles without considering local discontinuities. These discontinuities are often essential for coiled-coil function, as demonstrated in examples such as signal transduction (Ferris et al., 2012) and intracellular trafficking (Murray et al., 2016).

Recent years have seen a number of breakthroughs in protein structure prediction with the development of deep learning-based methods, the most prominent of which is AlphaFold2 (Jumper et al., 2021; Evans et al., 2022). It is implemented as an end-to-end sequence-to-structure model that also exploits evolutionary information provided in the form of a multiple sequence alignment. Benchmarks have demonstrated its superiority over classical homology modeling approaches (Lupas et al., 2021) and its applicability to the modeling of protein complexes (Akdel et al., 2022) and peptides (McDonald et al., 2023). In addition, although not designed for such tasks, AlphaFold2 has been reported to provide insight not only into protein structure but also into its dynamics (Winski et al., 2024; del Alamo et al., 2022; Stein and Mchaourab, 2022; Wayment-Steele et al., 2022). Finally, AlphaFold2 has also been used in the context of coiled coils, revealing, for example, that approximately 15% of protein-protein interfaces are mediated by these motifs (Schweke et al., 2024).

In this work, we present a systematic benchmark of modeling coiled-coil domains with AlphaFold2. Going into this project, there were features of coiled coils that we thought might be challenging for AlphaFold2. For example, experimental studies have shown that the topology and oligomerization state of coiled coils can be altered by one or a few mutations, typically near the hydrophobic core (Harbury et al., 1993; Yadav et al., 2006), suggesting that the alternative conformations may be separated by relatively low energy barriers. Regarding the use of evolutionary information, it is worth noting that the structural features and repetitive nature of coiled coils impose strong constraints on their sequences, which has been identified as a problem because it often leads to false matches between non-homologous coiled-coil sequences (Mistry et al., 2013). This could result in the MSA provided to AlphaFold2 being “contaminated” with non-homologous coiled-coil segments that could potentially have very different structural properties. With these potential challenges in mind, we set out to evaluate the ability of AlphaFold2 to predict coiled-coil features, including global topology and local geometry.

## Methods

### Datasets

The benchmark was to compare experimentally determined coiled-coil structures from CCdb (Szczepaniak et al., 2021) with their corresponding AlphaFold2 predictions. We focused on structures containing dimeric, trimeric, and tetrameric coiled-coil bundles, filtering out structures with resolutions below 3.5 Å or with complex topologies other than regular bundles. Finally, we retained only structures in which coiled-coil regions accounted for more than 50% of all residues to minimize the influence of non-CC regions on folding. All the filtering steps were performed using the localpdb package (Ludwiczak et al., 2022). This process resulted in an “automatic” benchmark set of 379 coiled-coil structures.

For the oligomer state prediction benchmark, we further refined this data set. We excluded heterooligomers and structures in which the heptad register could not be assigned with TWISTER (Strelkov and Burkhard, 2003) (unambiguous register assignment was critical because some of the benchmarked methods use this feature to improve prediction accuracy). This resulted in a set of 216 structures, with each oligomer class (38% dimers, 36% trimers, and 26% tetramers) similarly represented. In addition to these sets, we constructed two additional sets based on manually curated examples, consisting of 19 parallel (Szczepaniak et al., 2018) and 21 antiparallel (Szczepaniak et al., 2014) coiled-coil structures, most of which were GCN4 variants. The four benchmark sets are summarized in Supplementary Table 1, which also shows the identity to the sequence aligned with the lowest E-value (not higher than 1e-3) in the UniRef100 database.

### AlphaFold2 modeling

Modeling was performed with ColabFold (version 1.5.2, fb16be5c2d913ca9e4a4e779be82e16dc1b9edf1) (Mirdita et al., 2022) using the alphafold2_multimer_v3 model, max. 5 rounds of recycling, and no templates. The models were ranked according to the ‘multimer’ metric. The simulations were run in two modes, the first with multiple sequence alignments generated with ColabFold, and the second with the -- msa-mode single_sequence parameter to force single sequence predictions. It is important to note that although we modeled the entire sequences (i.e., including the non-coiled coil regions), we performed the structural analyses only on the coiled-coil regions.

### Oligomerization state prediction

We chose to benchmark LOGICOL (Vincent et al., 2013) and CoCoNat (Madeo et al., 2023) as representatives of state-of-the-art coiled-coil oligomerization state prediction methods. Unlike AlphaFold2, both programs can be run with coiled-coil register information, which improves the robustness of the predictions. We benchmarked LOGICOL and CoCoNat in two different scenarios, a) with the structurally derived heptad registers obtained with TWISTER (Strelkov and Burkhard, 2003), and b) without explicitly providing register information (indicated by the “noregister” suffix in Figure 2A). While CoCoNat implements its own register prediction, we opted to run LOGICOIL in the same way as its web server, with registers predicted by MARCOIL (Delorenzi and Speed, 2002).

**Figure 2.**
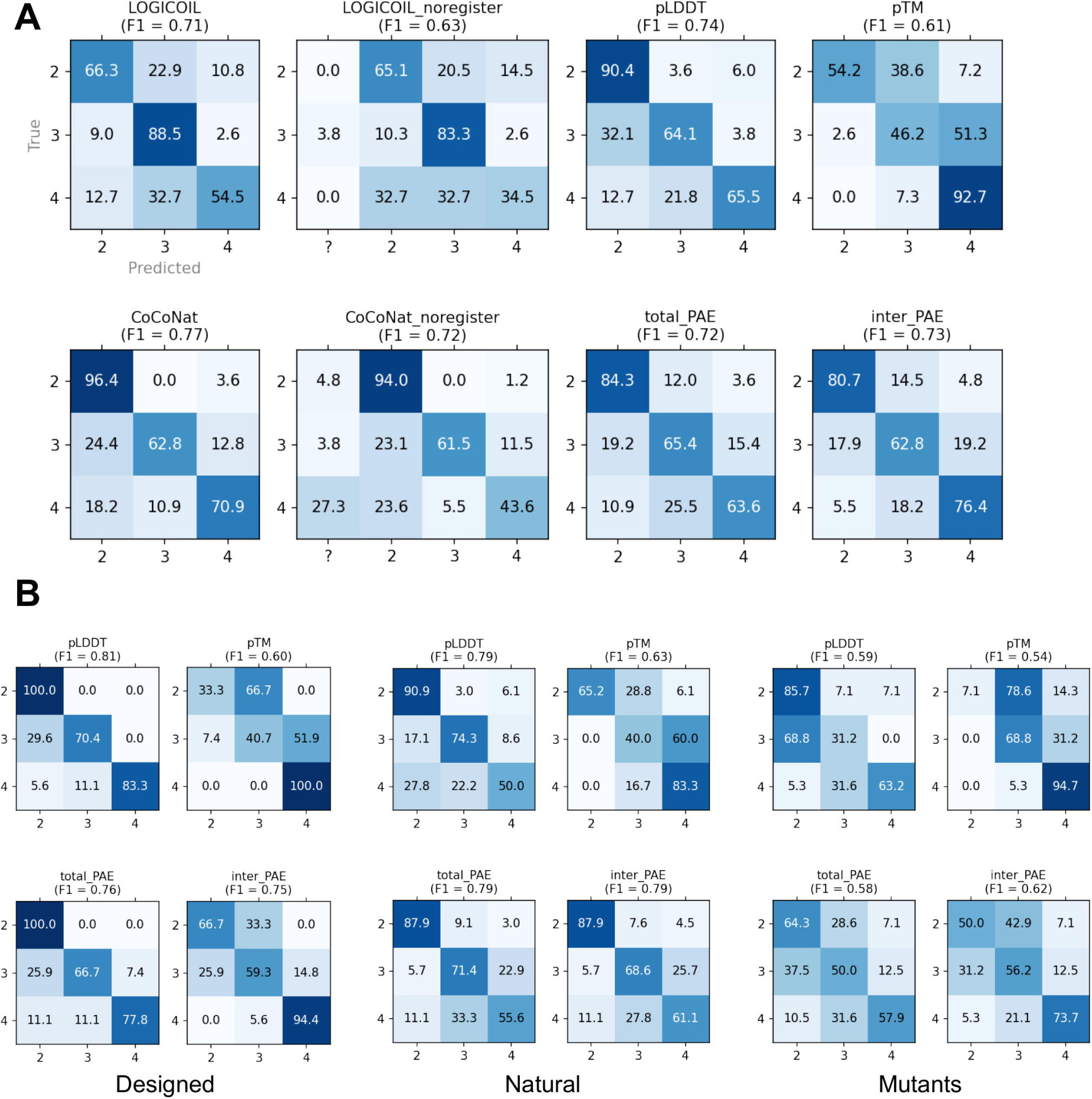
Accuracy of AlphaFold2, LOGICOIL and CoCoNat in predicting the oligomeric state of coiled coils. (A) LOGICOIL and CoCoNat were run in two modes, with and without (“_noregister” suffix) providing structurally derived heptad registers. AlphaFold2 predictions were made using the model quality metrics pLDDT, pTM and PAE, the latter divided into total_PAE and inter_PAE (see text for details). The accuracy of each method is shown as a confusion matrix with true and predicted labels on the y and x axes, respectively. Weighted F1 coefficients are also provided. (B) Confusion matrices computed for the subsets of the benchmark set corresponding to designed, mutant, and natural sequences.

To assess the extent to which AlphaFold2 can be used to predict the oligomeric state of coiled-coil domains, we used it to model each of the 216 sequences in the “oligomerization” benchmark set as a dimer, a trimer, and a tetramer, and checked whether the oligomeric state of the best-scoring model matched the oligomeric state observed in the experimental structures. We tested several metrics, including pLDDT, pTM, and PAE. We further refined the PAE score (total_PAE) into the interchain PAE (inter_PAE), which can be thought of as the uncertainty in predicting the relative position of the helices. Confusion matrices and weighted F1 scores were calculated for each method (Figure 2).

Given the good accuracy of the AlphaFold2 quality metrics in predicting oligomeric state, and the fact that AlphaFold2 was not directly supervised for oligomeric state prediction (Evans et al., 2022), we hypothesized that accuracy could be further improved by supervised learning on the internal representations of AlphaFold2. To test this, we extracted representations for each benchmark sequence modeled as a monomer (retrieved with the --save-single-representations option). The resulting representations were then averaged over the sequence length dimension, resulting in fixed size vectors of shape 5×256 (5 corresponds to each of the AlphaFold2 pre-trained models and 256 to the embedding size) per sequence. For visualization purposes, the dimensionality of the embeddings was reduced using PaCMAP (Wang et al., 2021) (Figure 4A, left panel).

To process these concatenated AlphaFold2 embeddings, we first trained a simple neural network using fivefold cross-validation. The neural network architecture included multiple dense layers with batch normalization, dropout, and L2 regularization. The model was trained using the Adam optimizer (Kingma and Ba, 2017) and a sparse categorical cross-entropy loss function. Early stopping with a patience of 3 epochs was applied during training to avoid overfitting. The dimensionality of the internal representations (from the last layer) of the model was reduced using PaCMAP (Wang et al., 2021) and visualized (Figure 4A, right panel). We also considered simpler classifiers available in the scikit-learn Python package, and based on fivefold cross-validation, we found LogisticRegression to be the best performing. The internal classifier parameters, namely regularization strength and class balance weighting, were optimized, and the final model, available on GitHub, was fitted to all available data without cross-validation (https://github.com/labstructbioinf/dc2_oligo).

### Bundle geometry prediction benchmark

We modeled each structure in the “automatic”, “parallel”, and “antiparallel” benchmark sets (see the Datasets section above) with the oligomerization state matching that of the experimental structure. For comparison with the corresponding experimental structures, we took the top-ranked model as representative of each prediction. We then superimposed the predicted and experimental structures using US-Align (Zhang et al., 2022), checking that all chains were aligned in the correct orientation and excluding misaligned cases from further analysis. Finally, we used SamCC Turbo to detect the coiled-coil segments and compute their structural parameters (helical axial rotation and supercoiling). An example of the comparison of these parameters between an experimental and a predicted model is shown in Figure 1C. Details on the calculation of these parameters can be found in (Szczepaniak et al., 2021).

## Results and Discussion

### Folding simulations and benchmark dataset

By default, AlphaFold2, as implemented in ColabFold, generates five independent models for a given sequence, each associated with its own set of quality scores. These scores are used to rank the models, with the top-ranked model (rank 1) being considered the best. This ranking system also applies to coiled-coil domains, where the ranks correlate with the structural similarity between the predicted and experimental structures (Supplementary Figure 1). However, it’s important to note that on average the differences between the ranks are small. Interestingly, AlphaFold2 model number 3 performed slightly better than the others.

To assess the quality of the AlphaFold2 models, we used 389 experimentally determined coiled-coil structures from the Protein Data Bank as a reference (Supplementary Table 1). The reference set is diverse, consisting of 147 dimers (100 parallel, 47 antiparallel), 115 trimers (101 parallel, 14 antiparallel), and 127 tetramers (53 parallel, 74 antiparallel). In addition, 61% of the structures are natural, showing 100% sequence identity to the most similar entry in the UniRef100 database, 24% are mutant variants (50%-99% identity), and the remaining 15% show no sequence similarity and are therefore considered de novo designs. In the subsequent analyses, each aimed at evaluating different aspects of coiled coil modeling, we used different subsets of the main benchmark set, namely “oligomerization”, “parallel”, “antiparallel”, and “automatic” (see Datasets section in Methods).

### Oligomerization state prediction

We began the benchmark by evaluating the performance of current methods for predicting the oligomerization state of coiled-coil domains, using the “oligomerization” set comprising 216 cases. In addition to the sequence-based methods LOGICOIL (Vincent et al., 2013) and CoCoNat (Madeo et al., 2023), we also evaluated the predictive power of the AlphaFold2 model quality metrics pLDDT, pTM, and PAE. For predictions based on AlphaFold2 scores, we computed models for oligomerization states ranging from dimer to tetramer for a given sequence. Then, for each metric, the oligomerization of a model with the best score was selected as the prediction. For each approach, we constructed a confusion matrix comparing the predicted number of helices with the number of helices in the experimental structures (Figure 2A). In addition, each confusion matrix is accompanied by a weighted F1 score, which is a quality metric ranging from 0 to 1. It combines precision and recall, giving more weight to classes that occur more frequently in the data. As expected, both LOGICOIL and CoCoNat performed well in the oligomerization state prediction task when provided with structurally derived coiled-coil registers, with F1 scores of 0.71 and 0.77, respectively, although LOGICOIL showed a strong bias toward predicting trimers, as did CoCoNat for dimers. When we did not provide the structurally derived heptad register (see Methods), the F1 scores dropped to 0.63 and 0.72, respectively (in addition to a larger number of incorrect predictions, some predictions were ambiguous, marked “?” in Figure 2A).

Among the AlphaFold2 metrics tested, the pLDDT score performed significantly better than pTM (F1 scores of 0.74 and 0.61, respectively), with the former showing a strong bias toward dimers and the latter toward tetramers. We also tested the PAE score, which is a 2D matrix representing the predicted error in the pairwise distances between all residues in all chains. In the benchmark, we considered the average of the entire PAE matrix (total_PAE) and the average of the PAE values between residues from different chains (inter_PAE). Both metrics performed similarly to pLDDT, achieving F1 scores of 0.72 and 0.73 for total_PAE and inter_PAE, respectively, but providing more balanced predictions across oligomerization states (Figure 2A). Predictions were then evaluated for the three subsets corresponding to de novo designs, natural sequences, and mutants (Figure 2B). While performance for designed and natural sequences was comparable to each other and to the full “oligomerization” set, predictions for mutants were less accurate (F1 scores for the pLDDT metric of 0.81, 0.79, and 0.59, respectively). Natural and designed sequences are better optimized for a given oligomerization state, either by natural selection or by rational design. In contrast, mutant sequences can often carry a conflicting signal, reflecting both the nature of their natural homologs and the specific functional or structural changes intended by the introduced mutations, which may explain the observed discrepancy.

Given the good accuracy of the oligomeric state predictions, we investigated how these predictions are reflected in the corresponding model quality metrics. The results show that most of the correct predictions (green dots in Figure 3A) are very close to the classification cut-off, i.e. the line separating them from the incorrect predictions (see dashed lines and histograms). This effect is particularly pronounced for adjacent oligomerization states (dimers vs. trimers and trimers vs. tetramers) and less noticeable when comparing dimers and tetramers, which are typically better separated (Figure 3B).

**Figure 3.**
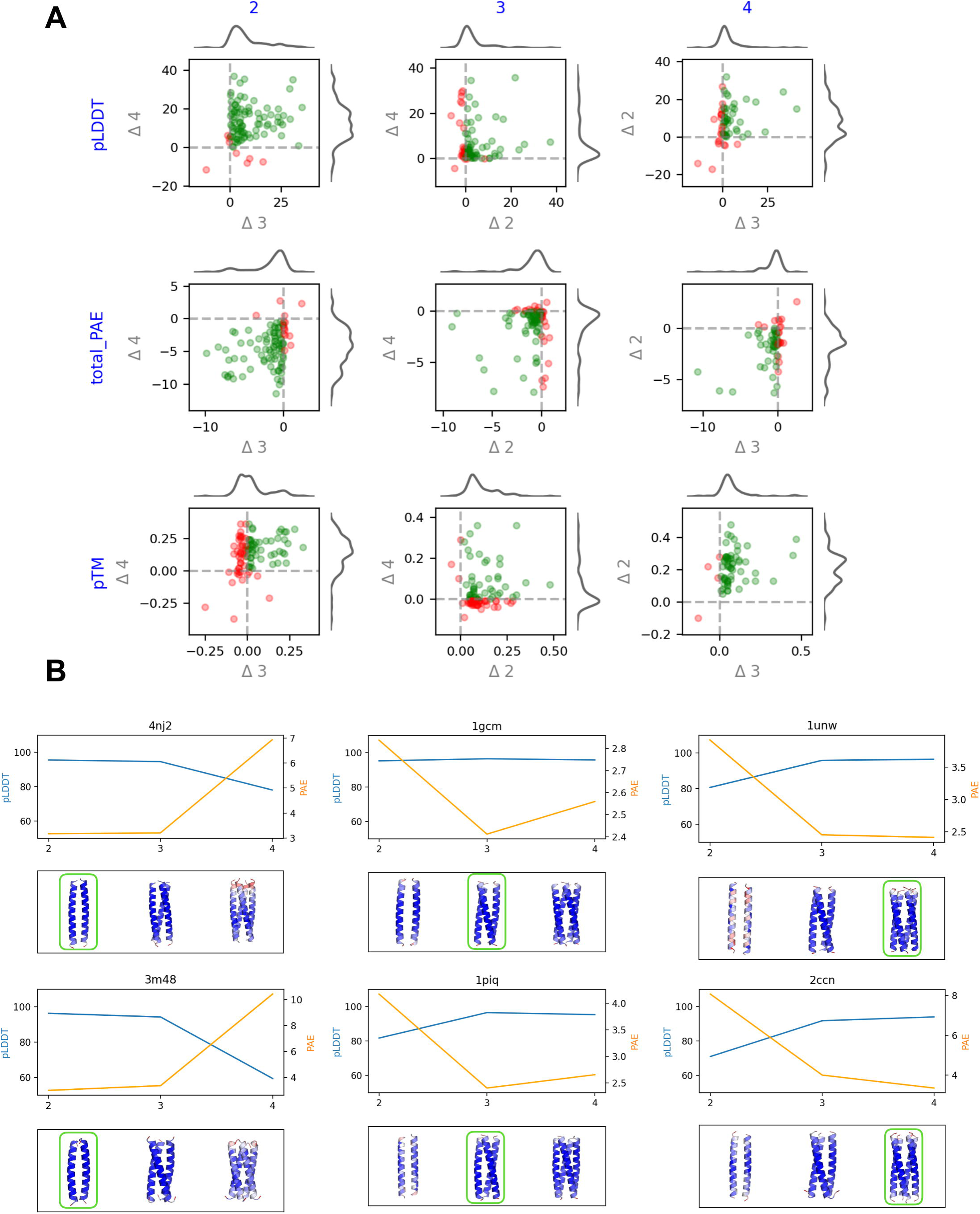
Accuracy of AlphaFold2 confidence scores in predicting oligomerization states of coiled-coil domains. (A) The columns represent the correct reference oligomerization states (dimers, trimers, and tetramers), while the rows correspond to the three AlphaFold2 quality scores: pLDDT, PAE, and pTM. Each scatter plot illustrates the difference between the score for the correct oligomerization state and the scores for the two incorrect states. Cases where the score for the correct state exceeds that for the incorrect states are shown in green. (B) pLDDT and PAE scores for selected benchmark cases modeled as dimers, trimers, and tetramers. The correct oligomerization state is highlighted with green boxes, and the structures are colored according to their pLDDT scores.

Finally, intrigued by the fact that AlphaFold2 had comparable accuracy to state-of-the-art methods despite not being trained to predict oligomerization state, we investigated whether such predictions could be made directly from the internal representations of AlphaFold2, rather than from the quality scores associated with models computed for different oligomerization states. These internal representations, also called embeddings, provide a numerical description of the input query sequence in the context of the knowledge captured by the AlphaFold2 model. They take the form of multidimensional vectors that can be compared to each other to reflect the similarity of the sequences they represent. The dimensionality of the representations obtained for multiple sequences can be jointly reduced using methods such as PaCMAP (see Methods for details), allowing them to be visualized as 2D maps that reflect AlphaFold2’s perspective on these sequences. Such a 2D map, obtained for all sequences from the “oligomerization” benchmark set, showed some degree of separation between dimers, trimers, and tetramers (Figure 4A, left panel). Motivated by this observation, we trained a simple neural network to predict a coiled-coil oligomeric state based on its corresponding AlphaFold2 non-reduced representation. Projection of the embeddings obtained from this neural network showed a clear separation between the groups (Figure 4A, right panel), suggesting the potential applicability of such an approach. In exploring alternative prediction models, we found that even a simple regression model can produce very good results: 5-fold cross-validation on the benchmark set showed significantly improved performance (F1=0.82) without any noticeable bias in the preferred oligomeric state (Figure 4B). Note, however, that we evaluated this model using 5-fold cross-validation, so the results obtained are not directly comparable to those shown in Figure 2. The implementation, along with the training routines, has been deposited on GitHub (https://github.com/labstructbioinf/dc2_oligo).

**Figure 4.**
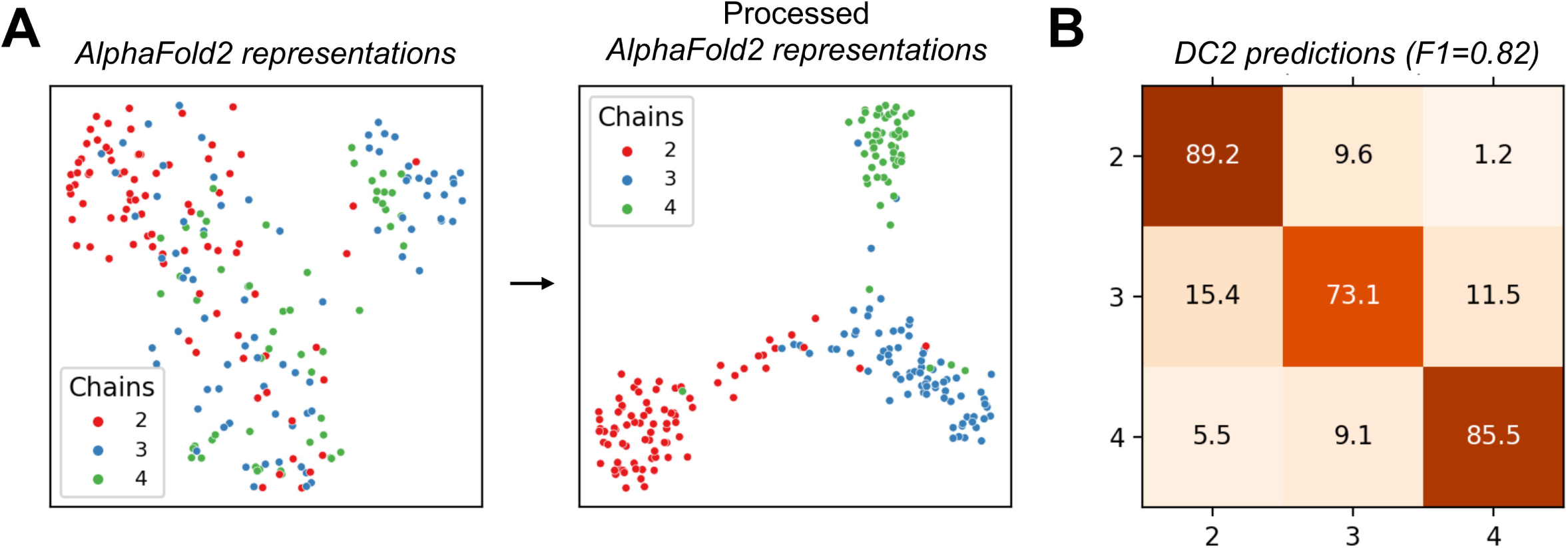
Application of AlphaFold2 representations for the prediction of the oligomeric state. (A) 2D plots of raw AlphaFold2 representations of the benchmark cases (left panel) and representations obtained from a downstream model (right panel). Colors represent oligomerization states. (B) A confusion matrix with true and predicted labels on the y and x axes, respectively, showing the performance of the final prediction model trained on AlphaFold2 representations. The weighted F1 score is also shown.

### Modeling of coiled-coil geometry

To perform a detailed comparison of the experimental structures with the corresponding AlphaFold2 models, we used SamCC (Szczepaniak et al., 2021), a program designed to automatically detect coiled-coil domains and calculate their structural parameters, including the degree of bundle supercoiling and hydrophobic core geometry (Figure 1). Comparisons were made using three separate sets of benchmarks, each designed to highlight a specific challenge in coiled-coil modeling. The first and second sets focused on packing geometries in parallel and antiparallel 4-helix bundles, respectively, while the third set was more general and included bundles with different oligomerization states and degrees of supercoiling. For each benchmark, all structures were modeled in the correct oligomerization state obtained from the corresponding experimental structure, and the models were then analyzed using SamCC.

The first set, termed “parallel”, consists of 19 canonical 7/2 parallel 4-helix bundles (13 of which are GCN4 variants) in which the helices interact by knobs-into-holes packing. Despite their overall similarity, these structures exhibit slight differences in the geometries of their hydrophobic cores. These differences are primarily manifested in the axial rotation of the helices, which is within ±5° (calculated relative to an idealized reference model of a coiled-coil employing knob-into-holes packing; Figure 1). Our previous work has shown that these subtle differences are not artifacts of the experimental procedures, but can be related to the sequence composition of the hydrophobic core (Szczepaniak et al., 2018). Among the structures analyzed, only a single GCN4 variant (1W5L), which exemplifies a rational design of an antiparallel to parallel structure, resulted in a model that incorrectly assumed an antiparallel topology. All other models obtained, including those for designed sequences such as 4H7R, showed the correct topology and their hydrophobic cores were modeled with high fidelity (Figure 5A).

**Figure 5.**
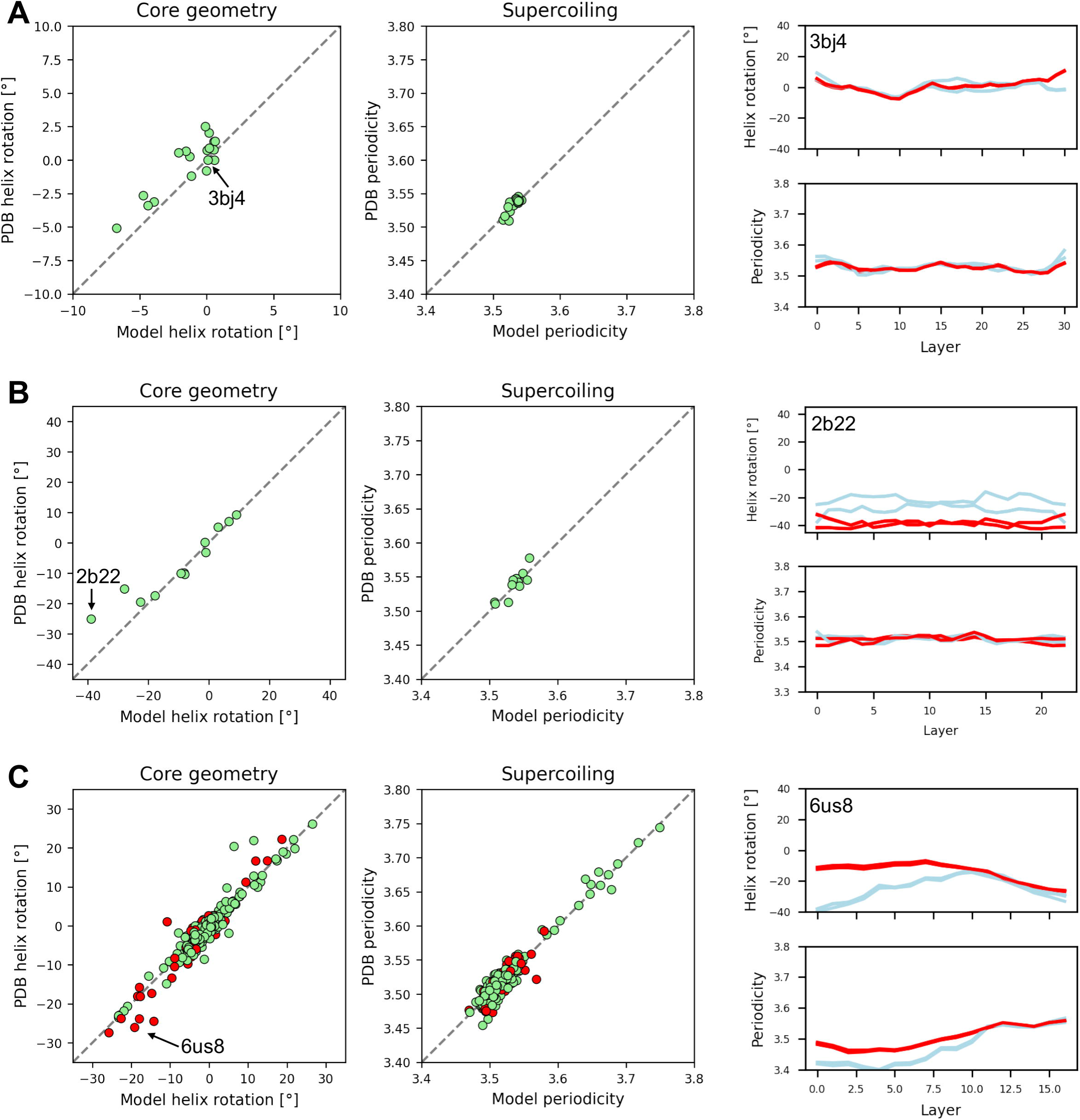
Comparison of coiled-coil parameters between experimental (PDB) and AlphaFold2 (model) structures. The scatter plots show the differences in average helix rotation and periodicity, while the accompanying plots (right) show detailed per-layer measurements for exemplary structures (model and experimental structures are shown in red and light blue, respectively). Panels A, B and C show the results for the “parallel”, “antiparallel”, and “automatic” benchmark sets, respectively.

The “antiparallel” set consists of 21 four-helix antiparallel bundles (9 of which are GCN4 variants) with hydrophobic core geometries defined by helix axial rotations ranging from -26° to +10°. Structures with rotations up to ±10° are considered to have canonical knobs-in-holes packing (Figure 1B, left panel), whereas those with rotations below -10° are considered to have non-canonical packing (Figure 1B, right panel). Canonical packing is characterized by the interaction of two residues per heptad repeat within the hydrophobic core. In contrast, non-canonical packing involves the co-option of an additional heptad position e into the core (Lupas et al., 2017). In this benchmark, AlphaFold2 correctly modeled only 12 out of 21 structures. In most of the unsuccessful cases, the resulting models had the wrong topology (parallel instead of antiparallel), preventing a meaningful comparison with the experimental structures. However, the remaining models with the correct topology, including those for de novo designed bundles such as 3S0R and 3TWE, were highly accurate in terms of core geometry (Figure 5B). The only exceptions were two coiled-coil domains (1ZV7 and 2B22), where co-opting the heptad position e to the core resulted in a strong negative rotation of the helices, which was underestimated in the models.

The two benchmark sets described above, i.e. “parallel” and “antiparallel”, contained hand-picked structures, mostly GCN4 variants. To evaluate AlphaFold2 more comprehensively, we generated a third “automatic” set containing 379 dimers, trimers, and tetramers of various topologies. Unlike the smaller benchmark sets, this set also included non-canonical bundles characterized by right-handed twist and periodicity >3.63 associated with repeats such as hendecads and pentadecads (Figure 1A). As with the two smaller sets, among the correct topology models (307 out of 379), the majority were accurate with respect to both helix packing and bundle periodicity (Figure 5C). Interestingly, the quality of modeling did not depend on the release date of the corresponding experimental structures, and those released after AlphaFold2 training (2018-04-30) were modeled equally well (compare red and green dots in Figure 5C).

### Bias in topology prediction

In both the oligomerization and geometry prediction benchmarks, AlphaFold2 showed a bias toward predicting parallel structures. In the oligomerization benchmark, the topology was correctly predicted for 93% of the parallel and 44% of the antiparallel bundles. In the geometric benchmark based on the “antiparallel” set, 9 out of 21 structures were incorrectly modeled as parallel. Given that eight of these are GCN4 variants, and that GCN4 WT adopts a parallel structure, it seems more plausible that this particular result is related to AlphaFold2’s inability to accurately model the effects of mutations, rather than a general bias toward parallel structures. The “parallel” set also contains many GCN4 variants, only one of which was incorrectly predicted to be antiparallel (GCN4-pLI E20C L16G variant; 1W5L). While GCN4-pLI (1GCL) is parallel, GCN4-pLI E20C (2CCN) is antiparallel, and the addition of the L16G mutation restores the structure to a parallel topology (1W5L). This example highlights the limitations of AlphaFold2 in predicting the effects of complex mutations, especially in the case of coiled coils, which can often adopt nearly isoenergetic alternative conformations.

Since the two sets discussed above consist primarily of GCN4 variants, we decided to use the entire benchmark set to estimate the accuracy of topology predictions and evaluate the associated quality scores. We found that AlphaFold2 performed well in predicting parallel structures, with only 10% incorrectly modeled as antiparallel. However, it showed a general bias in predicting antiparallel structures, with 42% modeled as parallel (Figure 6). A very similar pattern was observed when the data was divided into natural sequences, mutants, and de novo designs. We also found that the AlphaFold2 quality metrics (pLDDT, PAE, and pTM) did not clearly distinguish between correct and incorrect topologies, and that there was considerable overlap between the two (Supplementary Figure 2). Finally, we observed differences between the three sequence categories, with natural sequences showing a more pronounced separation between the scores associated with correct and incorrect predictions.

**Figure 6.**
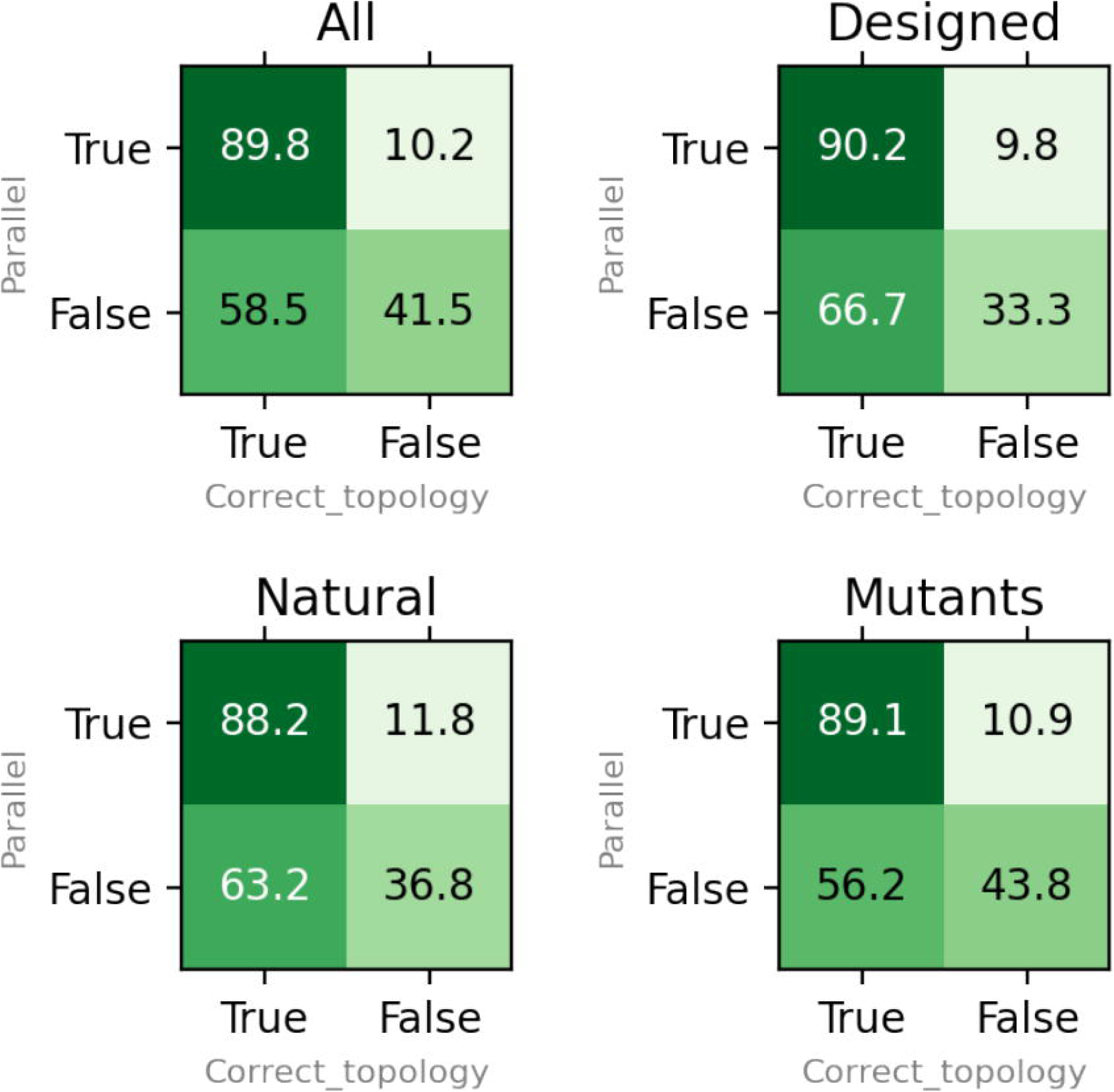
The matrices show the ratio of cases where the topology was correctly modeled when the correct oligomeric state was provided, categorized by helical orientation (parallel or antiparallel). Separate plots are shown for all benchmark cases (see Supplementary Table 1), as well as for dimers, trimers, and tetramers.

These results suggest that AlphaFold2 models are biased toward parallel structures; therefore, topology predictions should be treated with caution, even when accompanied by good quality metrics. We also observe a decrease in prediction quality for mutated and designed sequences; however, we note that correct predictions can still be achieved for rationally designed bundles, such as those included in our benchmark (3S0R and 3TWE) and studied by others (Naudin et al., 2022).

The effect of using multiple sequence alignments

By default, AlphaFold2 uses evolutionary information derived from a multiple sequence alignment (MSA), where the target sequence is aligned with its homologs. However, this approach has two potential drawbacks: first, the computation of MSAs is often time-consuming, sometimes taking longer than the actual folding process; second, some sequences have no detectable homologs, either as natural singletons or because they were designed de novo. To address these issues, we tested the performance of AlphaFold2 in modeling coiled-coil structures without the use of an MSA.

We first investigated how the removal of MSA information affected the prediction of the oligomeric state and, to our surprise, found no difference between the results with (Figure 2) and without (Supplementary Figure 3) MSA. We then evaluated the MSA-free mode in terms of topology prediction using the “parallel”, “antiparallel” and “automatic” benchmark sets described above. In the “parallel” set, all models predicted in the MSA-free mode had both correct topology and core-packing geometry, suggesting that MSA was not essential for these tasks. However, in the “antiparallel” set, we observed a drop in performance, with only 8 out of 21 cases having the correct topology in the MSA-free mode (compared to 11 cases when MSA was used). In addition, the quality of the packing geometry in some of these eight cases was poor, which was not a problem with the MSA-based models (Figure 5B).

To further understand the effect of MSA on topology prediction, we compared the AlphaFold2 confidence scores (pLDDT, total_PAE, and pTM) across the “automatic” set examples modeled with and without MSA. Specifically, we focused on 307 cases that were correctly modeled with MSA and used them as a reference point for the MSA-free modeling. In the majority of cases (>85%), we observed only small differences in scores between the MSA and MSA-free models (Figure 7). Furthermore, the MSA-free models had the correct topology in >95% of these cases. However, in the remaining cases where the MSA-free models had lower scores, many (>50%, shown in red in Figure 7) had incorrect topology. We also found that the average effective number of sequences in the MSAs was higher for the benchmark cases where the confidence scores were most affected by the lack of MSAs (174 vs. 48).

**Figure 7.**
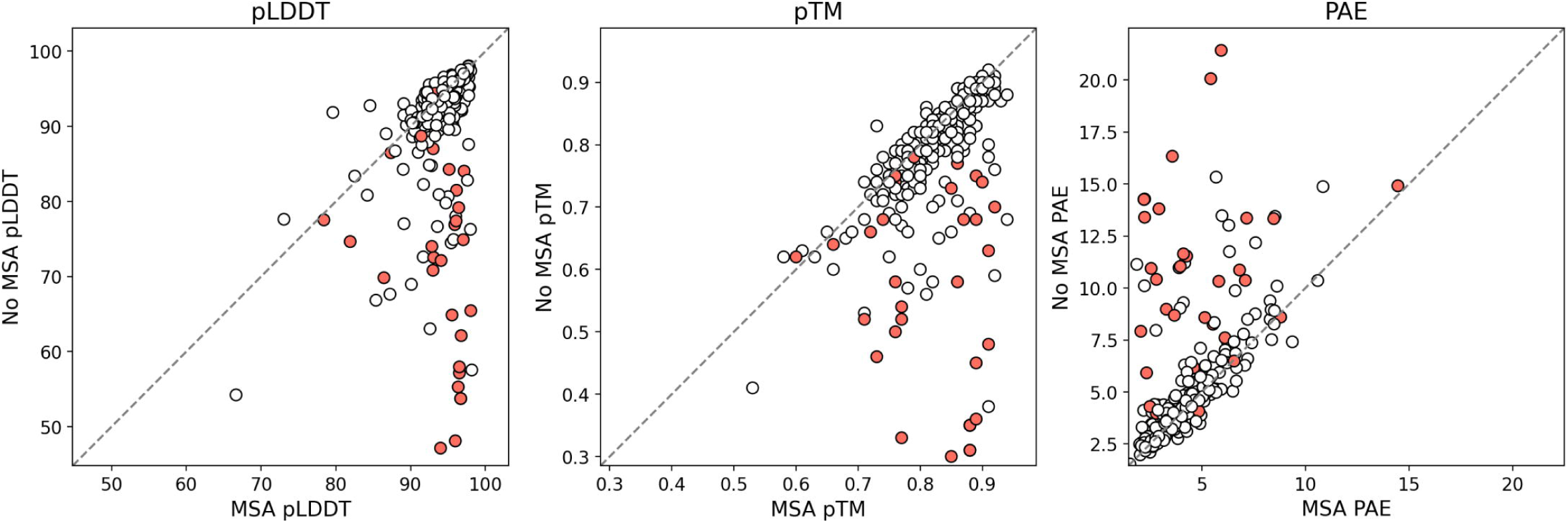
Effect of using multiple sequence alignments on AlphaFold2 quality metrics. Each scatter plot compares a specific AlphaFold2 quality metric (pLDDT, pTM, and PAE) associated with corresponding models calculated with (x-axis) and without (y-axis) the use of multiple sequence alignments. White dots represent benchmark cases where both MSA-guided and MSA-free modeling resulted in a model with correct topology, while red dots indicate cases where only MSA-guided modeling provided the correct topology.

These results show that the use of an MSA improves the accuracy of coiled-coil modeling, especially in cases where the MSA is rich in evolutionary information (i.e., has a high effective number of sequences). However, MSA-free modeling also produced accurate models in a significant number of cases, and topological errors could often be identified by their lower scores. Thus, the MSA-free mode may be useful for high-throughput screening, provided that strict score cutoffs are applied.

## Conclusions

While AlphaFold2 provides highly accurate predictions of oligomeric states (Figures 2, 3, and 4), as well as fine details such as supercoiling and core geometry (Figure 5), its ability to predict the correct orientation of helices, i.e., bundle topology, is limited (Figure 6). In all benchmarks, we observed a tendency for AlphaFold2 to model antiparallel structures as parallel. This problem becomes more pronounced when MSAs are not used during modeling (Figure 7), consistent with their critical role in the co-evolution-based prediction of long-range contacts that guide the proper orientation of secondary structure elements. For example, in the case of the human tetherin structure (3MQC), the modeling without MSA results in a misalignment of helices in the rank 1 model and even a switch to a parallel structure in the rank 2 model (Figure 8A). Interestingly, modeling with MSA restores the correct alignment of the helices, but the structure forms two separate dimers, which is not consistent with the experimental tetrameric form. The rank 2 MSA-based model, although tetrameric and more similar to the experimental structure, contains significant helical distortions that reduce the quality metrics. However, we note that the rank 1 model, although not matching the experimental tetramer, may still be more accurate in a functional sense, as the dimeric form of tetherin has been shown to be functional (Yang et al., 2010). Another example of the difference between the experimental structure and the corresponding models calculated with and without MSA is the human CtIP protein (7BGF, Figure 8B). Its rank 1 model calculated without MSA closely matches the experimental parallel structure; however, there is a clear discrepancy between the experimental structure and the AlphaFold2 rank 1 model obtained with MSA. While the experimental structure assumes a straight coiled coil, the AlphaFold2 MSA-based model appears to be broken in the middle. The observed break in the model results from the presence of a functionally important region, a zinc hinge, which is essential for the function of the protein (Morton et al., 2021). Such examples highlight the possibility that some of the AlphaFold2 models can provide clues to protein function even when they do not agree with experimental structures, and underscore the benefit of using MSA.

**Figure 8.**
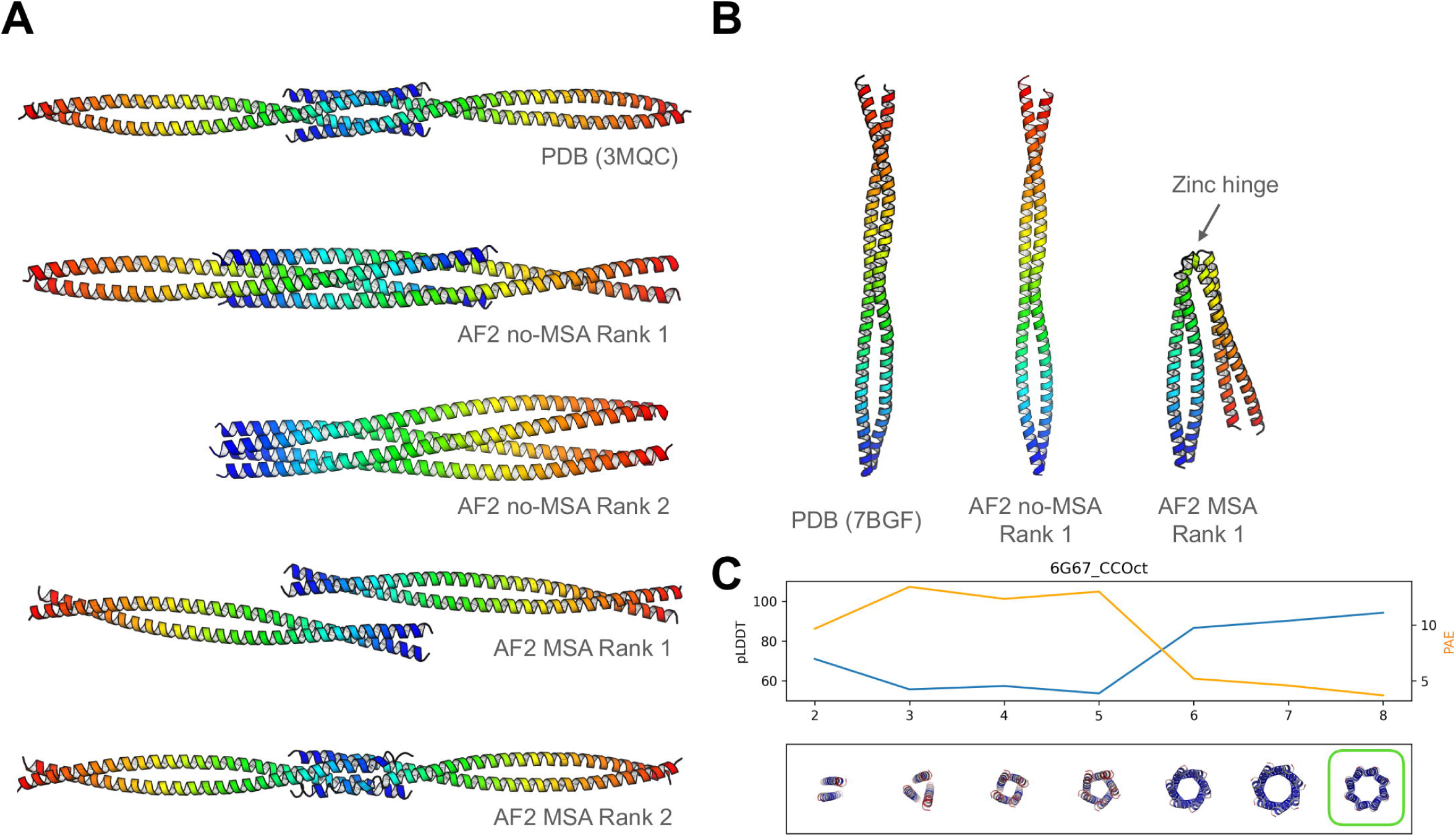
Examples of successes and failures in modeling coiled-coil structures with AlphaFold2. (A) modeling of tetherin and (B) CtIP proteins. See text for details. (C) pLDDT and PAE scores for a designed octameric coiled-coil barrel modeled from dimer to octamer. The correct oligomerization state is highlighted with a green box, and the structures are colored based on their pLDDT scores.

Surprisingly, the use of MSA has very little effect on the prediction of the oligomeric state. We propose that the success in predicting the oligomeric state in the MSA-free mode is due to the fact that in coiled coils it is determined by the presence of additional hydrophobic positions (Szczepaniak et al., 2014) or the formation of multiple interfaces by a single helix (Lupas et al., 2017). These features can be accurately described using only the target sequence and do not require evolutionary information. It is also worth highlighting the unexpected accuracy of predicting the oligomerization state using the internal AlphaFold2 representations. Such an approach proved to be not only effective (Figure 4), but also faster than modeling a given coiled-coil sequence in all possible oligomerization states (Figures 2 and 3). Recently, the AlphaFold2 embeddings have also been explored in the context of predicting protein-ligand interactions (Gazizov et al., 2023). Prediction of oligomerization states has also been attempted using ESM2 model embeddings (Sledzieski et al., 2023). We also note that new AlphaFold2-based methods for predicting structures of homooligomers (Schweke et al., 2024) and complex heteroligomeric assemblies (Shor and Schneidman-Duhovny, 2024) have recently been proposed.

Finally, although this work focused on bundle-shaped coiled coils, specifically those that adopt oligomeric states up to tetramers, we found that AlphaFold2 also shows potential for predicting the oligomeric states of coiled-coil barrels. In the example shown in Figure 8C, the correct oligomeric state, an octamer, is associated with the highest pLDDT and lowest PAE values. The applicability of AlphaFold2 to coiled-coil barrels will require further investigation.

## Acknowledgments

S.D-H. and M.M.G. were supported by institutional funds of the Max Planck Society. This work was also supported by the “Excellence Initiative - Research University” program, an internal grant of the University of Warsaw to enhance the research potential of its staff.

