## Supplementary Figure for "Applicability of AlphaFold2 in the modeling of dimeric, trimeric, and tetrameric coiled-coil domains"

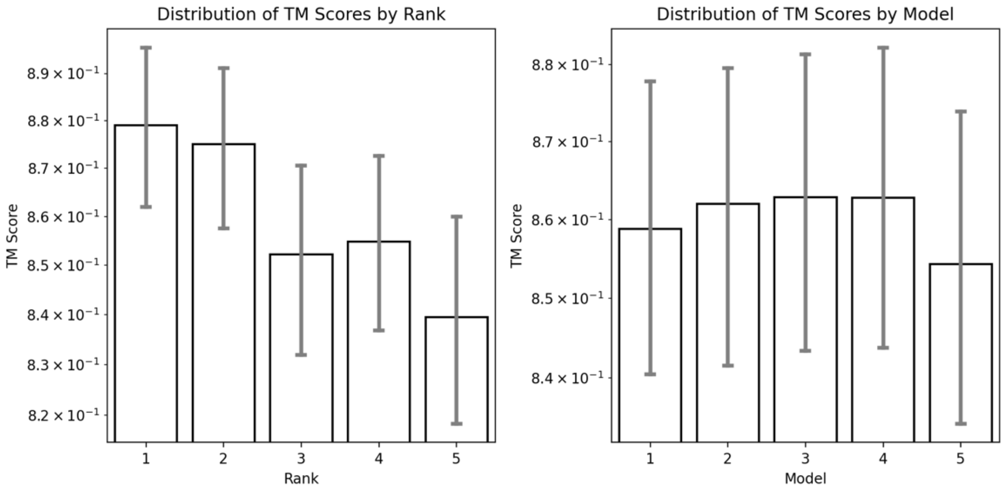


**Supplementary Figure 1.** Structural similarity between AlphaFold2 models and their corresponding experimental structures listed in Supplementary Table 1. TM score similarity metrics were calculated using US-Align run in oligomer mode (-mm 1 and -ter 0 parameters). The left panel shows the distribution of TM scores per AlphaFold2 rank, and the right panel shows the description of TM scores per AlphaFold2 model number. Note the logarithmic scale.

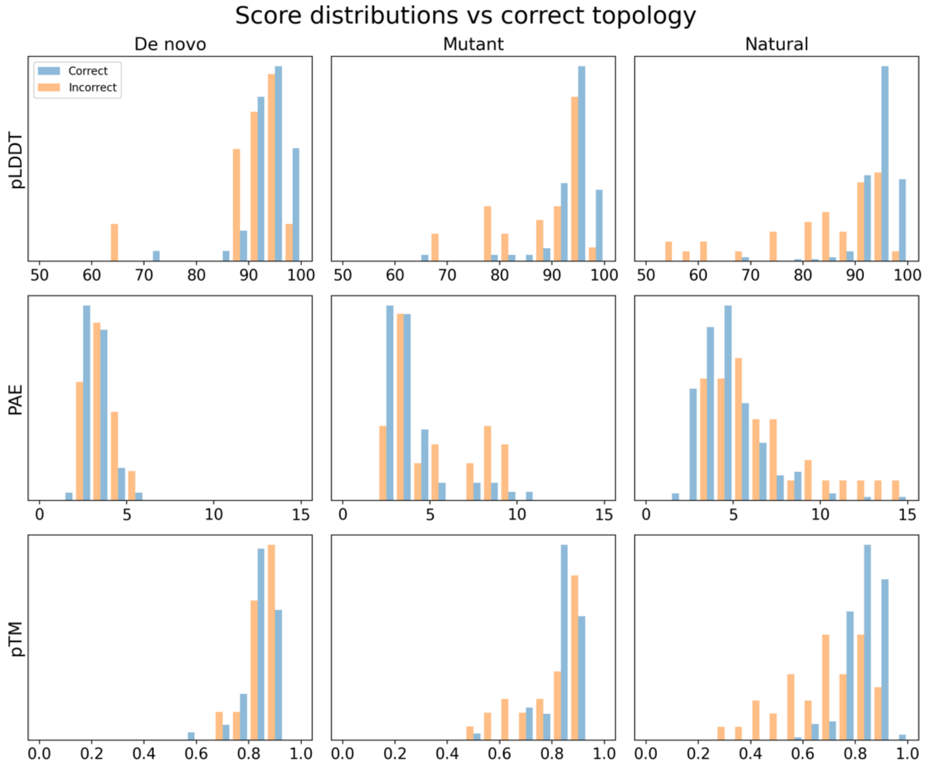


**Supplementary Figure 2.** AlphaFold2 quality metrics and topology prediction. In each plot, the scores associated with correct (blue) and incorrect (orange) predictions were normalized and plotted separately.

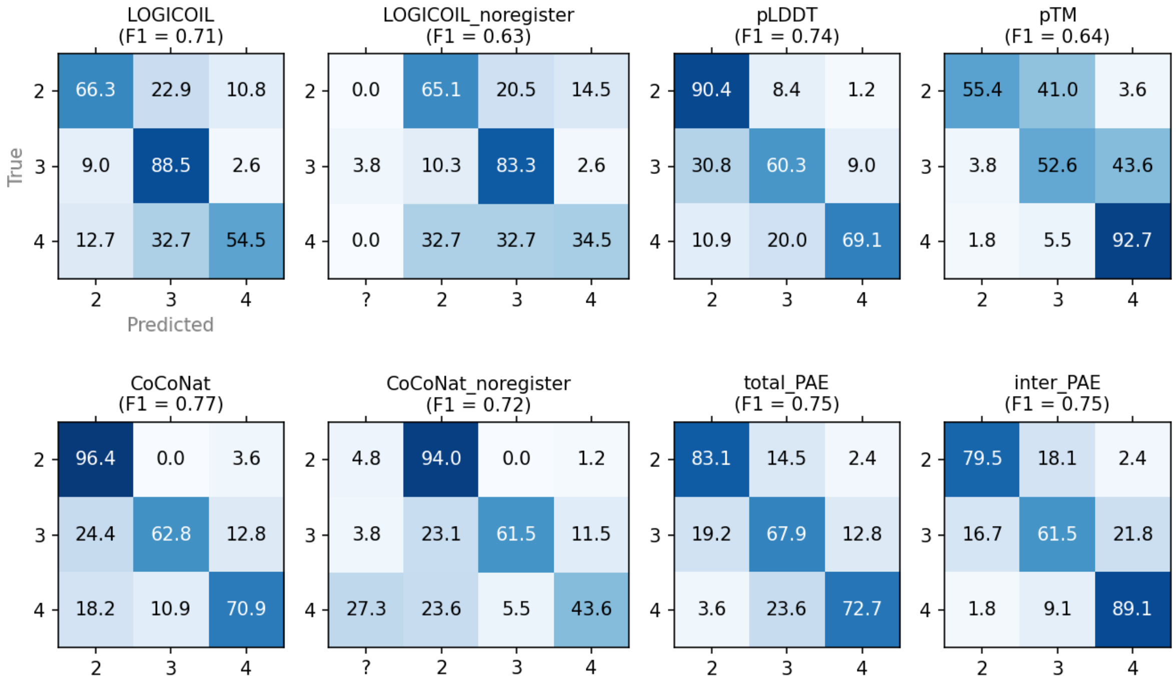


**Supplementary Figure 3.** Accuracy of AlphaFold2 run in no-MSA mode, LOGICOIL and CoCoNat in predicting the oligomeric state of coiled coils. See legend to Figure 2A in the main text for details.
